# The combination of optogenetic-induced protein aggregation and proximity biotinylation assays strongly implicates endolysosomal proteins in the early stages of α-synuclein aggregation

**DOI:** 10.1101/2024.10.16.618762

**Authors:** Maxime Teixeira, Razan Sheta, Dylan Musiol, Vetso Ranjakasoa, Jérémy Loehr, Jean-Phillippe Lambert, Abid Oueslati

## Abstract

Alpha-synuclein (α-syn) aggregation is a defining feature of Parkinson’s disease (PD) and related synucleinopathies. Despite significant research efforts focused on understanding α-syn aggregation mechanisms, the early stages of this process remain elusive, largely due to limitations in experimental tools that lack the temporal resolution to capture these dynamic events. Here, we introduce UltraID-LIPA, an innovative platform that combines the Light-Inducible Protein Aggregation (LIPA) system with the UltraID proximity-dependent biotinylation assay to identify α-syn-interacting proteins and uncover key mechanisms driving its oligomerization. UltraID-LIPA successfully identified 38 α-syn-interacting proteins, including both established and novel candidates, highlighting the accuracy and robustness of the approach. Notably, a strong interaction with endolysosomal and membrane-associated proteins was observed, supporting the hypothesis that interactions with membrane-bound organelles are pivotal in the early stages of α-syn aggregation. This powerful platform provides new insights into dynamic protein aggregation events, enhancing our understanding of synucleinopathies and other proteinopathies.

## Introduction

Alpha-synuclein (α-syn) aggregation into insoluble inclusions, known as Lewy bodies (LBs), is a hallmark of Parkinson’s disease (PD) and other synucleinopathies^1,2^. Since α-syn identification as the primary component of LBs^3^, its aggregation has been strongly linked to the neuronal loss characteristic of PD, positioning α-syn and its pathological forms as promising targets for disease-modifying therapies^4–7^.

This therapeutic potential of targeting α-syn has prompted extensive research into the mechanisms driving its aggregation and the effects of this process on neuronal homeostasis^8–11^. These studies have deepened our understanding of α-syn role in LB formation and toxicity, revealing insights into the composition, ultrastructure, and biochemical properties of these proteinaceous inclusions^12–14^. However, the majority of this research has focused on end-stage α- syn inclusions, leaving the early events of aggregation and the cellular machinery involved largely unexplored^15,16^. Moreover, the dynamic nature of α-syn aggregation requires experimental tools with high temporal resolution to capture early oligomerization steps and identify key proteins involved^17–20^.

To address these challenges, we present a novel approach combining two complementary methods. First, we utilize the Light-Inducible Protein Aggregation (LIPA) system, which generates α-syn inclusions that faithfully replicate key features of authentic LBs^21,22^. This system allows real-time induction and monitoring of α-syn aggregation in living cells and *in vivo* with unprecedented temporal accuracy^21^. Second, we use the UltraID-proximity-dependent biotinylation assay coupled with mass spectrometry, a powerful tool for probing protein-protein interactions in living cells^23^. This combination led to the development of the UltraID-LIPA platform, designed to identify proteins that interact with α-syn during its early conformational conversion from monomeric to aggregated forms, providing new insights into the molecular processes driving aggregation.

After validating the compatibility of the two methods in living cells, we used the UltraID-LIPA system and mass spectrometry analysis to identify a set of proteins that specifically interact with early α-syn aggregates. Strikingly, this set includes both previously known α-syn interactors, confirming the reliability of our approach, and newly identified proteins, highlighting the robustness of the method. Further validation through biochemical and immunocytochemistry techniques confirmed the physical interactions between α-syn aggregates and the chosen top hits, underscoring the accuracy of the approach.

An in-depth analysis of the data revealed that many of the identified interactors are membrane-related proteins, suggesting that cellular membranes play a crucial role in the early stages of α- syn aggregation. This observation aligns with clinical and experimental evidence showing that lipids and membranes are key constituents of LBs in both PD patients and cellular models of α- syn inclusion formation^12,13,24^.

In conclusion, we report a novel platform that leverages the temporal resolution of two state-of-the-art techniques, providing a powerful tool to study the α-syn interactome during its oligomerization course. This approach offers new insights into the dynamic cellular machinery involved in the early stages of protein aggregation and is versatile enough to be applied to any amyloidogenic protein, advancing our understanding of proteinopathies.

## Results

### Creation and characterization of UltraID-light-inducible protein aggregation (UltraID-LIPA) system to identify early α-syn aggregates interactome in living cells

Our exploration of the α-syn aggregates early interactome started by the creation of a construct combining an optimized version of the BioID system, referred to as UltraID, and LIPA-α-syn. To this end we fused the UltraID abortive biotin ligase to the N-terminal end of α-syn (UltraID-LIPA- α-syn) (**Fig. 1A**). UltraID-LIPA-α-syn construct was then stably expressed in Flp-in T-REx HEK293T cells under a doxycycline-inducible CMV promoter^25,26^. The UltraID system is a compact and hyperactive enzyme that allows for the biotinylation of proximal proteins in a ∼10 nm radius^23,27,28^ (**Fig. 1B**). The addition of exogenous biotin to the cells in culture upon exposure to the blue light will label proteins interacting with or in close proximity to α-syn during its structural conversion from monomeric to oligomeric forms (**Fig. 1B**). To ensure that analysis is specific to LIPA-α-syn, we used two control constructs, LIPA-Empty, a construct lacking α-syn, and LIPA fused to TAR DNA-binding protein-43 (LIPA-TDP-43), an aggregation-prone protein implicated in amyotrophic lateral sclerosis (ALS)^29,30^ (**Fig. 1A & C**).

**Figure 1.**
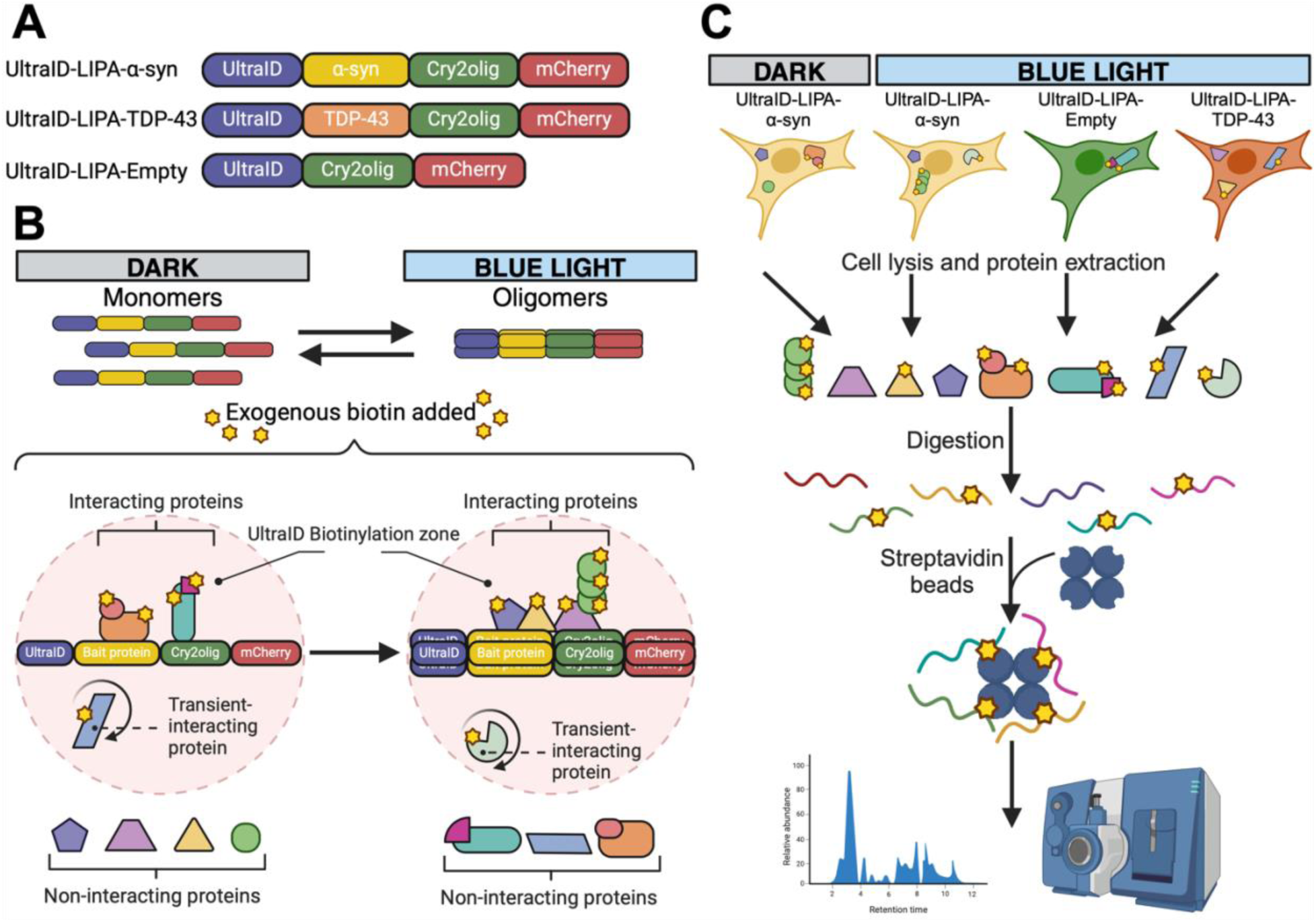
Description of the UltraID-LIPA constructs and the general workflow. (**A**) Schematic representation of the UltraID-LIPA constructs, composed of Cry2olig (green) fused to mCherry (red) at the C-terminal region. Wild-type α-syn (yellow) or TDP-43 (orange) were then fused at the N-terminal region of Cry2olig. Finally, UltraID was tagged to the N-terminal region of either α-syn (UltraID-LIPA-α-syn), TDP-43 (UltraID-LIPA-TDP-43) or directly to Cry2olig (UltraID-LIPA-Empty). (**B**) Schematic representation of the UltraID proximity biotinylating labelling. UltraID-fused constructs, expressed as monomers, undergo a rapid conversion to oligomers upon exposure to the blue light. After adding exogenous biotin, UltraID catalyses the biotinylating the proteins in contact or in close proximity (10 nm) with the bait protein. Most of the proteins will be transient or permanent interactors of the bait protein and are different depending on the status of the protein (either monomers or oligomers). (**C**) Schematic representation of the general workflow, from the cells to the LC/MS proteomic screening. The specific proteome of the newly formed α-syn oligomers is determined using several conditions: UltraID-LIPA-α-syn monomers (not exposed to blue light) are directly compared toUltraID-LIPA-α-syn oligomers formed under the blue light control. Other LIPA oligomeric constructs were used to control for the specificity of the α-syn interactome, namely UltraID-LIPA-Empty and UltraID-LIPA-TDP-43. The cells are then lysed, and the proteins are extracted using a RIPA lysis method combined with sonication. Extracted material was then digested, and the small peptides tagged with the biotin were captured using Streptavidin-Sepharose beads. The collected peptides were finally purified and analyzed using LC/MS.

First, we confirmed that fusing UltraID to α-syn does not impair the ability of LIPA-α-syn construct to aggregate and form LB-like inclusions in mammalian cells. As shown in Fig. 2A, cells overexpressing LIPA-α-syn form mCherry inclusions upon exposure to blue light (**Fig. 2A**). The size and fluorescence intensity of these inclusions increase positively with the duration of blue light exposure, consistent with our previous findings^21,31^ (**Fig. 2A**). A similar increase in the inclusions size and fluorescence intensity was also observed for the control constructs, LIPA- Empty and LIPA-TDP-43 (**Fig. S1**). Importantly, LIPA-α-syn, both with and without UltraID, exhibited a similar number of inclusions per cell at various light exposure times, indicating comparable aggregation propensities for the two constructs (**Fig 2B**). Moreover, the fusion of UltraID did not affect the phosphorylation of LIPA-α-syn inclusions at Ser129 (pS129), a key biochemical feature of pathological LBs^32–34^ (**Fig. 2C**). Collectively, these data confirm that the addition of the UltraID sequence to the LIPA construct does not influence LIPA-α-syn aggregation propensity or the key events involved in this process.

**Figure 2.**
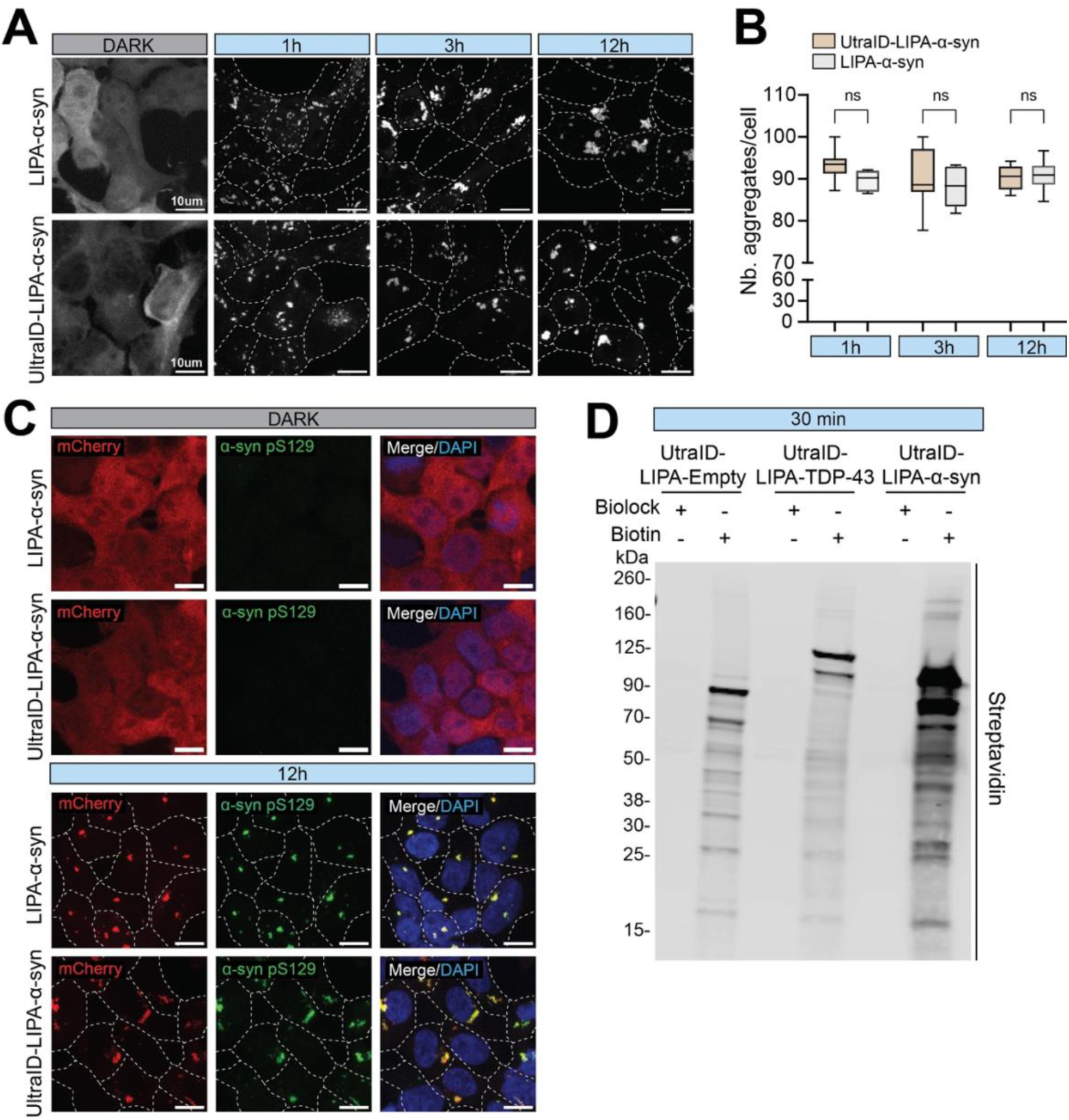
UltraID-LIPA-α-syn aggregates biotinylates proximal proteins while displaying authentic features of α-syn inclusions. (**A**) Confocal maximum intensity projections images of Flp-In T-REx HEK293T cells stably expressing UltraID-LIPA-α-syn construct. Cells were exposed to blue light for 1h, 3h and 12h to induce the aggregation. Scale bar = 10 µm. (**B**) Box plots graphs showing a quantification comparing the cells stably expressing the UltraID-tagged aggregates and the non-tagged, evaluating the number of cells bearing aggregates after several timepoints of illumination. Statistical differences were assessed with a two-Way ANOVA test, comparing the non-tagged cells and the tagged-cells overtime (ns = not significant). (**D**) Confocal maximum intensity projections images of HEk cells overexpressing LIPA-α-syn or UltraID-LIPA-α-syn, exposed (12h) or no to the blue light and stained for the pathological α-syn phosphorylation at the residue S129 (pS129). In both conditions we were able to detect pS129 staining after 12h of blue light exposure. Scale bar = 10 µm. (**D**) Representative Western blot membrane from Flp-In T-REx HEK293T cells stably expressing UltraID constructs, probed for Streptavidin. Cells were cultured with Biolock in the culture medium to block any endogenous biotinylation. For the biotinylation assay, biolock was removed and exogenous biotin was added at 50 µM to the cells whenever they were also exposed to blue light, for 30 min. Note that Streptavidin shows positive staining only in the conditions where exogenous biotin was added for all the UltraID constructs, with a signal accumulated at the expected size of the constructs (∼90/125/110 kDA, for UltraID-LIPA-Empty, UltraID-LIPA-α-syn and UltraID-LIPA-TDP-43, respectively). (

Finally, we assessed whether UltraID biotinylation capacity could be affected by the clustering of the LIPA constructs. After 24 h of doxycycline treatment, which provided optimal expression levels of UltraID-LIPA-α-syn, UltraID-LIPA-Empty, and UltraID-LIPA-TDP-43 constructs in the cells, we added exogenous biotin to the media and exposed the cells to blue light for 30 minutes to induce first steps of light-induced α-syn clustering. Western blot analysis of the biotinylation activity of each UltraID-LIPA construct revealed significant accumulation of biotin-labeled proteins, in contrast to the control conditions incubated with Biolock, a biotinylation-blocking agent (**Fig. 2D**). This observation confirms the capacity of the UltraID-LIPA system to label endogenous proteins under an aggregation process. Collectively, our data confirmed the compatibility of the LIPA and UltraID approaches for studying the protein interactome during the aggregation process. Furthermore, a 30-minute exposure to blue light and exogenous biotin is established as a compatible time point to study the α-syn interactome during the early stages of its aggregation process.

### Proximity biotinylation assay identified membrane-related proteins as the major interactome during α-syn early oligomerization steps

Of the 684 proteins screened in our proximity dependent biotinylation assay (**Table Sup 1**), 117 were found to interact with monomeric α-syn (no blue light), while 567 were more specifically associated with oligomeric α-syn forms (30 min blue light; **Fig. S2, volcano plot**). We then employed the SAINTexpress algorithm^35^ to filter out proteins non-specifically interacting with α- syn aggregates, comparing them to those interacting with monomeric LIPA-α-syn (no blue light) and LIPA-TDP-43 aggregates. This additional filtering allowed us to refine our results further, leading to the identification of 38 specific proteins that were significantly enriched (false discovery rate (FDR) ≤ 1%) with α-syn early inclusions (**Fig. 3A**). Among the identified proteins, a GO:term analysis revealed that 18 α-syn aggregate-interacting proteins were enriched in the endolysosomal (**Fig. 3A, yellow and green backgrounds**) and membrane-related compartments (**Fig. 3A, green and blue backgrounds**). We also identified 9 proteins that were linked to Ubiquitin-conjugation (**Fig. 3A, purple background**), (**Fig. 3A**). Finally, we were able to associate 7 proteins with genomic regulation, with an additional 3 associated with histone regulation (**Fig. 3A, pink and dark brown**), 4 proteins defined as RNA-binding (**Fig. 3A, dark and light brown backgrounds**) and 6 associated with acetylation (**Fig. 3A, bold characters**). Other proteins (10) were not associated with any specific cellular pathway (**Fig. 3A, white background**).

**Figure 3.**
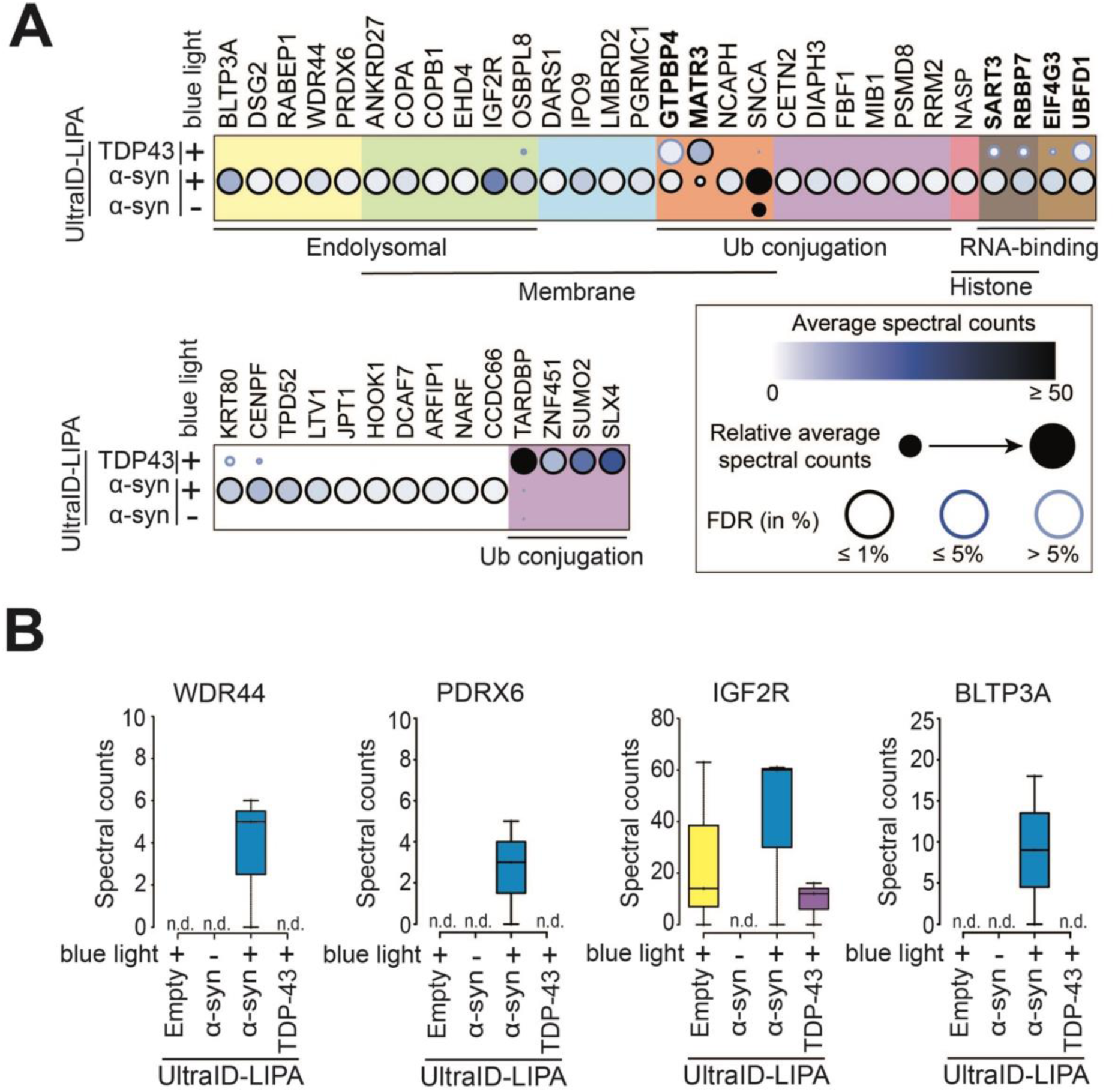
UltraID-LIPA system identifies key proteins intracting with α-syn early aggregates. (**A**) Dot plots of selected interaction partners associated with LIPA-α-syn monomers (not exposed to blue light), LIPA-α-syn early aggregates (exposed to blue light), or LIPA-TDP-43 (exposed to blue light). Circles are pseudo-colored to represent the average spectral counts for each of the markers represented. Each circle is also magnified depending on their relative average spectral counts, and an outer circle is pseudo-colored to represent the percentage of False Discovering Rate (FDR). The hits were arranged using a GO-term cellular compartment analysis. (**B**) Raw spectral counts are displayed for key relevant hits selected to be specific interactors of LIPA-α-syn oligomers, belonging to the endolysosomal family: WDR44, PRDX6, IGF2R and BLTP3A.

It is important to note that some of the identified protein hits have been previously described as direct α-syn interactors (*e.g.*, RABEP1^36^, COPB1^36^, OSBPL8^36^, PRDX6 (NSGP)^37^, PGRMC1^38,39^, DARS^36^, KRT80^38,39^, EIF4G^36^ and PSMD2/3^39^) or as potential modulators of α-syn aggregation (IGF2R^40,41^) or α-syn prion-like propagation (COPB1^42^). Meanwhile, other proteins were identified for the first time as α-syn interactors, using our approach, (BLTP3A, DSG2, WDR44, ANKRD27, COPA, EHD4, DARS1, IPO9, LMBRD2, GTPBP4, MATR3, NCAPH, CETN2, DIAPH3, FBF1, MIB1, RRM2, NASP, SART3, RBBP7, UBFD1, CENPF, TPD52, LTV1, JPT1, HOOK1, DCAF7, ARFIP1, NARF and CCDC66). Similarly, analysis of LIPA-TDP-43 interactome confirmed 2 known hits related to TDP-43 aggregation (ZNF451^43^, SUMO2^43–46^) out of 3 significant proteins (FDR ≤ 1%) identified in the study. This observation indicates that the approach is both reliable and robust, allowing for subtle or previously overlooked interactions with α-syn aggregates to be identified.

Moreover, detailed spectral counts analysis for four of the most specific hits, including known α- syn interactors (IGF2R, PRDX6) and newly identified ones (WDR44, BLTP3A), revealed that despite biological variation between samples, certain interactions occur exclusively with aggregated forms of α-syn (WDR44, PRDX6, and BLTP3A). This suggests that these interactions might be specifically driven by the pathological conformational changes of α-syn to the oligomeric state. On the other hand, we identified a candidate (IGF2R) capable of interacting with other LIPA aggregates, suggesting that this protein might have an affinity to the oligomeric conformation regardless of the aggregable protein present, however, it still showed enhanced affinity for aggregated α-syn (**Fig. 3B**).

### Validation of top interactors hit using biochemical and immunocytochemistry assays

Next, we sought to confirm the physical interaction of our candidates of interests (IGF2R, WDR44, PRDX6 and BLTP3A) with α-syn aggregates using co-immunoprecipitation, thus validating the robustness of the proteomic data. The experimental design consists of exposing HEK293T cells stably expressing LIPA-α-syn (without UltraID) to the blue light for 1 h, followed by 30 min of blue light stimulation in the presence of the crosslinker disuccinimidyl glutarate (DSG) in order to stabilize the protein complexes (**Fig 4A**). DSG is an irreversible amine-reactive crosslinker, with a short space arm of 7.7 Å and commonly used in studying α-syn protein-protein interaction with protein partners as well as stabilizing its structure^47–52^.

**Figure 4.**
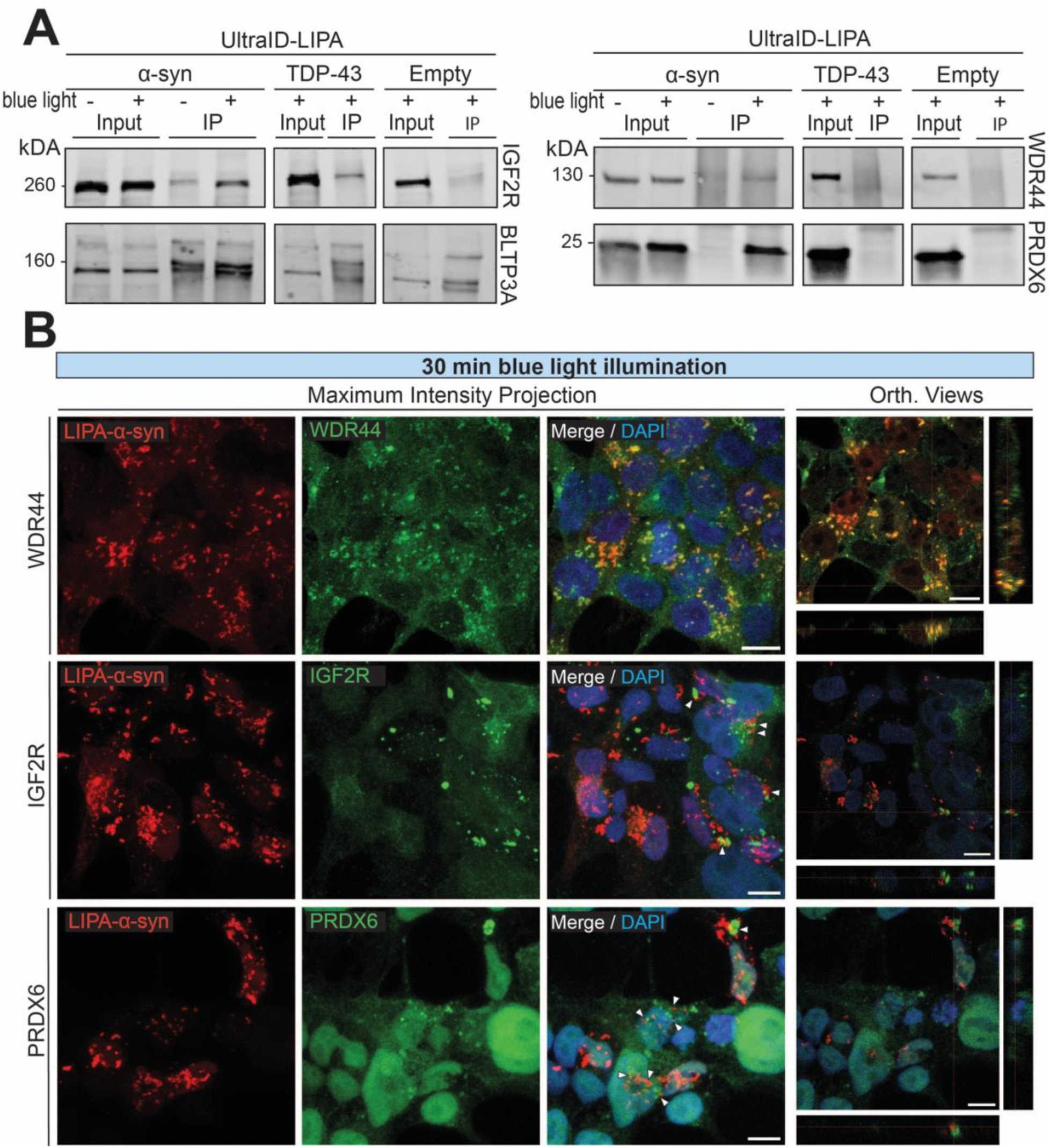
Validation of the specificity of the proteomic hits. (**A**) Representative nitrocellulose membranes showing the immunoprecipitated material. LIPA-α-syn, LIPA-TDP-43 and LIPA-Empty crosslinked aggregates were immunoprecipitated from HEK293T cells exposed to blue light illumination at 0.3 mW/mm^2^ for 90 min. WDR44 and PRDX6 were only detected in the LIPA-α-syn condition exposed to blue light. IGF2R and BLTP3A were minimally detected under control conditions but were expressed at significantly higher levels in the LIPA-α-syn condition upon blue light exposure. Uncropped membranes are available on Fig. S3 (**B**) Confocal maximum intensity projections images (left) and orthogonal views (right) of HEK293T cells stably expressing the LIPA-α-syn construct. Cells were exposed to blue light for 30 min to induce aggregation and were fixed for immunocytochemistry. Note that α-syn aggregates are highly colocalizing with WDR44, as shown by the orange color in the merged image, and with the orthogonal views (right panel). PRDX6 and IGF2R are accumulating next to the aggregates, making direct contact with them (merged image, white arrows; orthogonal views). Scale bar = 10 µm.

We used magnetic beads coupled with an antibody targeting mCherry to pull down α-syn aggregates and then probed the Western blot membrane for the four hits identified in our proteomic screening. The choice of mCherry was made to prevent potential issues such as epitope loss or reduced antibody affinity caused by conformational changes in α-syn when using anti-α-syn antibodies, as well as being able to use the same antibody to pull down the control LIPA constructs. Interestingly, Western blot revealed selective and enhanced enrichment of the proteins of interest (WDR44, IGF2R, PRDX6 and BLTP3A) in the presence of α-syn aggregates, in comparison with immunopurified LIPA-α-syn monomers (- blue light), LIPA-TDP43 and LIPA- Empty aggregates (**Fig. 4A**) (**Fig. S3**, uncropped Western blots).

Finally, we used immunocytochemistry to further confirm the interaction of α-syn aggregates with the proteins of interest. Of note, this approach allows, thanks to its spatial resolution, to appreciate complex formation and to discriminate if two proteins are either colocalizing or are close proximity partners. Interestingly, we could detect after 30 min of blue light stimulation, a convincing colocalization of WDR44 and LIPA-α-syn aggregates (**Fig 4B, top panel**), while PRDX6 and IGF2R are showing an accumulation at the vicinity and in contact with α-syn aggregates (**Fig. 4B, middle and bottom panel**).

Collectively, our findings validate the efficiency and relevance of combining proximity biotinylation assay and the LIPA system to identify key proteins implicated or interacting with α-syn during the early stages of its pathological conformational structural change and oligomerization. These data also emphasize the possible important role of proteins associated to the endolysosomal system in α-syn aggregation in living cells.

## Discussion

Protein aggregation is a complex and dynamic process that remains poorly understood, largely due to the limitations of current experimental models. To address this limitation, we need an approach combining a sensitive detection of protein proximity with a system allowing for the controlled initiation of α-syn aggregation with a tight temporal resolution in living cells. In this study, we combined the UltraID-proximity-dependent biotinylation assay and the LIPA system to capture, in real-time, an unprecedently studied concept of early α-syn aggregation stages in living cells. Our investigation started by validating the combination of the two approaches, fusing an UltraID moiety, that play a role in catalysing biotin attachment to the proximal proteins, with the LIPA-α-syn system, and data showed that this approach was successful as the combinations of the two constructs did not affect the aggregation kinetics nor the key biochemical events associated to the aggregation process, namely α-syn phosphorylation at Ser129 (pS129). Moreover, the UltraID fusion remained active during the blue light treatment and maintained its biotinylating activity as α-syn underwent conformational structural changes and oligomerization.

Using the UltraID-LIPA system, we identified proteins interacting with or in close proximity to α- syn during the early events of its oligomerization. Several controls were employed to minimize potential artifacts, allowing us to refine our analysis and protein identification. Notably, we confirmed multiple previously known α-syn interactors (RABEP1^36,53^, COPB1^36^, OSBPL8^36,54^, PRDX6 (NSGP)^37^, PGRMC1^38,39,55^, DARS^36^ and EIF4G^36,56^) and aggregation modulators (IGF2R^40,41,57–59^ and COPB1^42^), validating the accuracy of our approach.

While the UltraID-LIPA system is a powerful tool, enabling us to explore early α-syn aggregation interactome, it might present some limitations. First, the obtained results will need to be validated using native α-syn constructs in the context of neuronal culture and/or in human tissue. Additionally, it is possible that fusing the UltraID biotin abortive ligase to the N-terminus, and the CRY2-mCherry to the C-terminus of α-syn may interfere with normal protein interactions, despite our results that suggest that the UltraID fusion does not impact aggregation. Repositioning all fused proteins to either the N-terminus or C-terminus, rather than both, could help minimize interference with protein partners and enable the identification of new α-syn interactors.

In-depth analysis of our data revealed that the majority of α-syn-interacting proteins were associated with the endolysosomal system and intracellular membranes, two major pathways identified to be enriched in purified LBs from PD patients^60,61^. Using a proximity dependent biotinylation proteomic screening, through either antibody recognition or APEX2, the vesicular pathways (exosomes, synaptic vesicle, endocytic trafficking) were found to be directly interacting with α-syn in either tissue from PD patients or in iPSC-derived neurons, emphasizing the relevance of our results^36,39^. Our results also tend to indicate that other pathways are also directly involved with α-syn oligomers, namely the ubiquitin-proteasome pathway, histone-binding, RNA- modulation, and acetylation. Those pathways were also emphasized by multiple groups, either as directly interacting with α-syn or directly found in LBs from patients^36,61–63^. Diving into the specific hits found in our study, we have reported the presence of the protein PRDX6 that plays a crucial role in oxidative stress mediation^64,65^. PRDX6 has been found to be increased in the brain of PD patients^37^ and other PRDXs, like PRDX3 and PRDX5 have been reported to be found directly in Lewy bodies and suggested as potential direct interactors of α-syn oligomers^60,61^, as well as localized to the lysosomal compartment^66,67^. Proximity dependent biotinylation assays, directly in primary neurons, have also reported some of the hits detected in our study as directly interacting with α-syn: EIF4G, RABEP1, COPB1, and OSBPL8^36^. Other proteins like PGRMC1, KRT80, ARF1 (other subunit distinct from ARF3) or PSMD2/3 (other subunits distinct from PSMD8) were also detected using biotinylation by antibody recognition, targeting the pS129 α- syn directly in PD tissues^39^. Finally, other proteins were reported to be linked to PD, but not reported as direct interactors of α-syn: In addition, we discovered novel direct interactors of α-syn that have been previously linked to PD through whole exome sequencing, gene expression analysis or transcriptomic analysis (BLTP3A^68,69^, NCAPH^70^, MIB1^71^, PSMD8^70,72^, RRM2^73^, SART3^70,74^, UBFD1^75^, CENPF^76,77^, LTV1^78^, JPT1^79^, HOOK1^70^) as well as linked to changes in protein levels in PD patients or models (WDR44^80^, ANKRD27^81,82^, PSMD8^83,84^, RRM2^85^, RBBP7^86,87^). This is strengthening our findings, emphasizing the fact that α-syn might interact with very specific proteins while transitioning from a monomer to an oligomer, questioning the role of the proteins non yet linked to PD in our dataset that are linked to the endolysosomal system like DSG2^88,89^, COPA^90^ or EHD4^91–93^. In addition, the specific interactors we have identified here might be different from the proteins found in mature Lewy bodies, as the interactome has been shown to be evolving over the maturation time of the aggregation thus explaining the differences in some of the hits detected^12^. It might be interesting to further investigate the evolving interactome using our LIPA system, comparing the hits detected at the early timepoints with a later timepoint of illumination (12h to 24h of blue light stimulation) to further compare what are the proteins interacting with soluble oligomers and more mature and insoluble oligomers.

In conclusion, our study highlights that the UltraID-LIPA system combines two powerful techniques: the biotin proximity proteomic assay and the optogenetic-based LIPA system. This innovative integration holds great promise for uncovering proteins and cellular pathways involved in the initiation and progression of protein aggregation, which is critical in PD and other proteinopathies. Additionally, the system’s unique temporal resolution provides a valuable tool for addressing unresolved questions regarding the kinetics, dynamics, and evolution of the complex and enigmatic process of protein aggregation.

## Acknowledgments

This work was supported by the Canadian Institutes of Health Research (CIHR) to A.O., the Natural Sciences and Engineering Research Council (NSERC; RGPIN-2023-05581) to A.O (NSERC; RGPIN-2024-04260) to J.P.L. and Society Parkinson Canada to A.O. A.O. and J.P.L. were supported by Junior 2 salary Awards from the Fonds de Recherche du Québec – Santé (FRQS) and la Société Parkinson du Québec (A.O.). M.T. was supported by scholarships from the Fondation du CHU de Québec (Bourse d’excellence Didier-Mouginot), FRQS and Parkinson- Québec Chaudière Appalaches. We thank the Proteomics Platform of the CHU de Québec Research Center for processing the sample.

Fig. 1 Created in BioRender. Oueslati, A. (2024) BioRender.com/y40f436

## Author contributions

Conceptualization: M.T., A.O.; Data curation: M.T., J.P.L.; Formal analysis: M.T., J.P.L., A.O.; Funding acquisition: A.O.; Investigation: M.T., R.S., D.M., V.R.; Methodology: M.T., J.L., J.P.L., A.O.; Project administration: A.O., M.T.; Resources: J.L., J.P.L., A.O.; Supervision: A.O., M.T.; Validation: M.T., A.O., J.P.L.; Visualization: M.T., R.S., J.P.L., A.O.; Writing-original draft, review, and editing: M.T., R.S., A.O., J.P.L.

## STAR Methods

### Key Resources Table

| REAGENT or RESOURCE | SOURCE | IDENTIFIER |
| --- | --- | --- |
| **Antibodies** |  |  |
| α-syn pS129 | Abcam | ab51253 |
| Alexa Fluor 488 | Invitrogen | A11008 |
| IGF2R | Proteintech | 20253-1-AP |
| UHRF1BP1 (BLTP3A) | Proteintech | 25121-1-AP |
| VPS13C | Proteintech | 28676-1-AP |
| WDR44 | Bethyl Laboratories | A301-440A |
| **Chemical, Peptides, and Recombinant proteins** |  |  |
| Bovine serum albumin (BSA) | BioShop | Cat#ALB001.500 |
| DAPI (4',6-diamidino-2-phenylindole) | Thermo Fisher Scientific | Cat#1729803 |
| D-Biotin | Bio Basic | BB0078 |
| Doxycycline | Takara | 631311 |
| BioLock | IBA LifeSciences | 2-0205-050 |
| Normal goat serum | Thermo Scientific | Cat. # 16210064) |
| Paraformaldehyde (PFA), EM grade | Electron Microscopy Sciences | Cat#19210 |
| Poly-L-lysine | ScienCell | 0413 |
| **Experimental models: Organisms/Strains** |  |  |
| HEK293T | ATCC | CRL-3216 |
| HEK293T-Rex | ThermoFisher | R78007 |
| **DNA Plasmids** |  |  |
| pcDNA-CMV Cry2olig-mCherry (LIPA-empty) | Dr. Abid Oueslati | N/A |
| pcDNA-CMV α-syn-Cry2olig-mCherry (LIPA-α-syn) | Dr. Abid Oueslati | N/A |
| pcDNA-CMV TDP-43-Cry2olig-mCherry (LIPA-TDP-43) | Dr. Abid Oueslati | N/A |
| pFRT-TO UltraID-3xFlag -LIPA-empty | Dr. Abid Oueslati | N/A |
| pFRT-TO UltraID-3xFlag-LIPA-α-syn | Dr. Abid Oueslati | N/A |
| pFRT-TO UltraID-3xFlag-LIPA-TDP-43 | Dr. Abid Oueslati | N/A |
| pOG44 | ThermoFisher | V600520 |

### Contact for Reagent and Resource Sharing

Further information and requests for resources and reagents should be directed to the Lead Contact, Dr. Abid Oueslati

### Method details

#### Cloning

pcDNA CMV templates of Cry2olig-mCherry (LIPA-empty) and α-syn-Cry2olig-mCherry (LIPA-α- syn) were generated and described in Bérard et al.^21^. pcDNA CMV template of TDP-43-Cry2olig- mCherry (LIPA-TDP-43) have been generated replacing α-syn from the LIPA-α-syn pcDNA CMV template, by the TDP-43 gene from pcDNA template wild type TDP-43-tdTOMATO-HA (Addgene, Cat. 28205)^94^. XhoI-HF (NEB, Cat. R0146S) and KpnI-HF (NEB, Cat. R3142S) restriction enzymes have been used according to manufacturer’s instructions to cut out the α-syn and TDP- 43 from their respective template. Digested material has been purified through 1% agarose gel, and the corresponding band for the TDP-43 insert and pcDNA CMV template lacking α-syn (TDP- 43: 1.2 kb; empty template: 6.2 kb) have been purified using PureLink™ Quick Gel Extraction and PCR Purification kit (Thermofisher, Cat. K220001). Purified material has then been ligated together using T4 DNA ligase (NEB, Cat. M0202T), following a ratio of 3:1 (insert:vector). The ligated material was finally transformed in DH5-α competent cells (NEB, Cat. C2987H) and amplified using GenElute™ HP Plasmid Maxiprep Kit (Sigma, Cat. NA0310-1KT).

To generate UltraID constructs, PCR amplification of the ORFs from the pcDNA templates (Cry2olig-mCherry, α-syn-Cry2olig-mCherry and TDP-43-Cry2olig-mCherry) were made using mutagenic primers flanking the ORFs with attB1 and attB2. Briefly, forward primers were designed as follows to match a N-ter tag within the same reading frame: 5’-GGGG ACA AGT TTG TAC AAA AAA GCA GGC TTC (25 gene specific nucleotides)-3’. Reverse primers were designed as follows: 5’-GGG GAC CAC TTT GTA CAA GAA AGC TGG GTC CTA (25 gene specific nucleotides)-3’. PCR products were then allowed to migrate through a 1% agarose gel, and the corresponding band for each insert was purified using PureLink™ Quick Gel Extraction and PCR Purification kit (Thermofisher, Cat. K220001). Purified PCR products were then subcloned into a Gateway-compatible pDONR223 vector following manufacturer’s instructions by site-specific BP recombination using BP Clonase™ II Enzyme mix (Thermofisher, Cat. 11789020) with an overnight incubation with the enzyme mix at 25°C. Obtained pEntry were transformed in DH5-α competent cells (NEB, Cat. C2987H) and amplified using QIAprep Spin Miniprep Kit (Qiagen, Cat. 27104). Generated entry vectors were then cloned into a gateway-compatible destination vector pDEST Flp-In UltraID-3xFlag-5’ by performing site-specific LR recombination following manufacturer’s instructions using LR Clonase™ II Enzyme mix (Thermofisher, Cat. 11791020) with an overnight incubation at 25°C. Resulting lentiviral vectors were transformed using One Shot™ Stbl3™ Chemically Competent cells (Thermofisher, Cat. C737303) and amplified using GenElute™ HP Plasmid Maxiprep Kit (Sigma, Cat. NA0310-1KT).

pDEST Flp-In UltraID-3xFLAG-5’ was generated by amplifying the UltraID-3xFLAG insert from pSF3 UltraID vector (Addgene, Cat. 172878)^23^ using mutagenic primers to induce NheI (before the insert; primer: 5’-ATA TGC TAG CAT GTT CAA GAA CCT GAT CTG GCT G-3’) and NsiI (after the insert; primer: 5’-ATA TAT GCA TCT TCT CCT TGA ACT TCT TCA GG-3’) cleavage sites. Cleaved material with NheI-HF (NEB, Cat. R3131S) and NSiI (NEB, Cat. R0127S) was then purified as previously described and ligated inside a pDEST FRT-TO TurboID^95^. Resulting pFRT- TO UltraID constructs were then transformed in One Shot™ Stbl3™ Chemically Competent cells (Thermofisher, Cat. C737303) and amplified using GenElute™ HP Plasmid Maxiprep Kit (Sigma, Cat. NA0310-1KT).

Lentiviral pcDNA-CMV were made for each construct (LIPA-empty, LIPA-α-syn and LIPA-TDP- 43), cloning pEntry previously generated into a Gateway-compatible destination vector pLenti CMV Puro DEST (w118-1) (Addgene, Cat. 17452) using the previously described LR reaction.

### Cell culture and generation of stable cell lines

HEK293T (ATCC, Cat: CRL-3216) and Flp-In T-REx HEK293T cells (Thermofisher, Cat. R78007) were maintained in high-glucose Dulbecco’s modified Eagle’s medium (DMEM) (Sigma-Aldrich, Cat. D5706) supplemented with 10% fetal bovine serum (FBS) (Sigma-Aldrich, Cat. F1051) and 1% penicillin/streptomycin (ThermoFisher, Cat. 15-140-122) at 37°C and 5% CO_2_.

Cell lines stably expressing UltraID constructs were made using the Flp-In™ T-REx™ system. Briefly, Flp-In T-REx HEK293T cells were seeded in a 6-well plate to reach 80% confluency the day of transfection. Cells were transfected with a standard calcium phosphate approach, using 200 ng of the plasmid UltraID FRT-TO and 1.8 µg of the expression vector pOG44 (Thermofisher, Cat. V600520). The next day (day 2), the cells were trypsinized and replated into 10-cm plates and selected with 200 µg/ml of hygromycin B as early as day 3. The selection media was then changed, every 3 days, until visible clones were obtained (approximately 2-3 weeks). Cells were then expanded and used for proteomic screening.

HEK293T cells stably expressing the constructs lacking the UltraID were made using lentiviral particles generated from pLenti DNA templates corresponding to each construct. Lentiviral particles were produced in HEK293T cells transfected with the templates in combination with packaging plasmids from 3^rd^ generation systems, following the protocol from Tiscornia *et al*.^96^. Lentiviral particles were then purified from the cell culture medium and titrated using a qPCR Lentivirus Titer Kit (ABM, Cat. LV900). HEK293T cells were then infected with the viral particles at a multiplicity of infection (MOI) of 5. Infected cells were subjected to selection pressure using puromycin at 1.5 µg/ml, and positive colonies were manually picked to obtain a homogenous mCherry expression.

### Biotin labeling of protein interactors and blue light exposure

Cells stably expressing UltraID constructs (UltraID-LIPA-empty, UltraID-LIPA-α-syn, UltraID- LIPA-TDP-43) were seeded in 15-cm plates at 2x10^6^ cells per plate in DMEM 10% FBS 1% P/S supplemented with BioLock (IBA Life Sciences, Cat. 2-0205-050) diluted at 1000X to minimize endogenous biotinylation. After 48h, doxycycline (Takara, Cat. 631311) was added to the medium to reach 1 µg/µL for 24 h to induce the expression of the constructs. The next day, cells were rinsed twice with warm dPBS Ca+ Mg+ (Gibco, Cat. 14040117) to remove remaining BioLock and complete medium supplemented with 50 µM of biotin (BioBasic, Cat. BB0078) was added to the cells. Cells that do not require blue light illumination (negative control for monomeric screening) were placed back in the incubator at 37°C and 5% CO_2_. Cells that required a blue light illumination (0.3 mW/mm^2^) were placed in an incubator equipped with a blue light system as described previously^21,22^. After 30 min incubation with the biotin medium, the reaction was stopped placing the plates on ice, removing the medium, and washing the cells once with ice-cold dPBS Ca+ Mg+. All further steps were performed on ice. The cells were scraped in 1 mL of cold PBS and collected in 2 mL collection tubes by centrifugation at 500 x g for 1 min at 4 °C. Cell pellets were snap frozen on dry ice and stored at −80 °C until streptavidin purification.

### Streptavidin beads purification

Cell pellets were thawed on ice and lysed in 1.5 mL of RIPA lysis buffer (1% NP-40, 0.1% SDS, 50 mM Tris-HCl pH 7.4, 75 mM NaCl, 0.5% Sodium Deoxycholate, 1 mM EDTA, 0.1 mM PMSF, 1 mM DTT, 1x Protease/Phosphatase inhibitor cocktail). All subsequent steps were performed on ice. Samples were sonicated for 30 s at 35% amplitude 10 s ON and 2 s OFF with a Q125 sonicator (QSonica, Cat. Q125-110). 250 Units of turbonuclease (Sigma, Cat. T4330-50KU) was then added to each sample and further incubated on a nutator for 1 h at 4 °C. Cells were centrifugated at 12,000 x g for 20 min at 4 °C. Supernatants were collected in a 2 ml tube and coupled with Streptavidin-Sepharose beads (Cytivia, Cat. 17511301) previously washed 2 times with RIPA lysis buffer. After an incubation of 3 h with rotation at 4 °C, the mix of sample-beads was spun down using a table-top centrifuge at max speed for 10 s. The beads were then washed with 1 mL of SDS-wash buffer (25 mM Tris-HCl, pH 7.4, 2% SDS, 10% Glycerol) to remove non-specifically bound proteins. After a spin down, beads were transferred to a clean 1.5 ml tube and resuspended in RIPA lysis buffer lacking PMSF and Protease/Phosphatase inhibitor cocktail. After a spin down of 10 s at max speed, the beads were washed 3 times with 1 mL of 50 mM ammonium bicarbonate (Bio Basic, Cat. AB0032). After the last wash, the beads were resuspended in 100 µl of 50 mM ammonium bicarbonate supplemented with 1 µg of MS-grade trypsin (Millipore Sigma, Cat. T6567) and incubated overnight at 37°C with rotation. The next morning, another 0.5 µg of trypsin was added to each sample and the digestion was resumed at 37°C for another 3 h. The beads were then pelleted by centrifugation for 2 min at 1,000 x g and the supernatant (containing the digested peptides linked to the beads) was transferred to a clean 1.5 ml tube. The remaining beads were washed 2 times with 200 µl of MS-grade 100% acetonitrile (Fisher Chemicals, Cat. A955-1) and the washed material was combined with the digested peptides previously collected. The pooled material was acidified by adding formic acid (Fisher Chemicals, Cat. A117-50) to the solution to a final concentration of 2% to quench the digestion. The samples were then dried in a SpeedVac centrifugal evaporator at 30°C for ∼2 h, and the dried peptides were then desalted using homemade C_18_ StageTips as demonstrated in Rappsilber *et al*.^97^. The final samples were then stored at -80°C until analysis by mass spectrometry.

### Mass spectrometry acquisition using an Orbitrap Fusion mass spectrometer

Peptide samples were separated by online reversed-phase nanoscale capillary liquid chromatography and analyzed by electrospray MS/MS. The experiments were performed with a Dionex UltiMate 3000 RSLCnano chromatography system (Thermo Fisher Scientific) connected to an Orbitrap Fusion mass spectrometer (Thermo Fisher Scientific) equipped with a nanoelectrospray ion source. Peptides were trapped at 20 μL/min in loading solvent (2% acetonitrile, 0.05% TFA) on an Acclaim 5 μm PepMap 300 μ-Precolumns Cartridge Column (Thermo Fisher Scientific) for 5 min. Then, the precolumn was switched online with a laboratory-made 50 cm × 75 μm internal diameter separation column packed with ReproSil-Pur C_18_-AQ 3- μm resin (Dr. Maisch HPLC) and the peptides were eluted with a linear gradient of 5–40% solvent B (A: 0,1% formic acid, B: 80% acetonitrile, 0.1% formic acid) over 90 min at 300 nL/min. Mass spectra were acquired in DDA mode using Thermo XCalibur software version 3.0.63. Full scan mass spectra (350–1,800 m/z) were acquired in the Orbitrap using an AGC target of 4e5, a maximum injection time of 50 ms, and a resolution of 120,000. Internal calibration using lock mass on the m/z 445.12003 siloxane ion was used. Each mass spectrometry (MS) scan was followed by MS/MS scans of the 10 most intense ions for a total cycle time of 3 s (top speed mode). The selected ions were isolated using the quadrupole analyzer in 1.6 m/z windows and fragmented by higher energy collision-induced dissociation at 35% collision energy. The resulting fragments were detected by the linear ion trap in rapid scan rate with an AGC target of 1e4 and a maximum injection time of 50 ms. Dynamic exclusion of previously fragmented peptides was set for 20 s and a tolerance of 10 ppm.

### Mass spectrometry data analysis

MS data was stored, searched, and analyzed using the ProHits laboratory information management system^98^. Thermo Fisher Scientific RAW MS files were converted to mzML and mzXML using ProteoWizard (version 3.0.4468^99^). The mzML and mzXML files were then searched using Mascot (version 2.3.02) and Comet (version 2012.02 rev.0) against the RefSeq database (version 57, January 30th, 2013) acquired from NCBI, which contains 72,482 human and adenovirus sequences supplemented with common contaminants from the Max Planck Institute (http://141.61.102.106:8080/share.cgi?ssid=0f2gfuB) and the Global Proteome Machine (GPM; http://www.thegpm.org/crap/index.html). Charges of +2, +3, and +4 were allowed, the parent mass tolerance was 12 ppm, and the fragment bin tolerance was 0.6 amu. Deamidated asparagine and glutamine and oxidized methionine were allowed as variable modifications. The results from each search engine were analyzed through TPP (the Trans-Proteomic Pipeline (version 4.6 OCCUPY rev 3^100^) via the iProphet pipeline^101^. Two unique peptide ions and a minimum iProphet probability of 0.95 were required for protein identification. SAINTexpress version 3.6.3^35^ was used to calculate the statistical probability of each potential protein-protein interaction compared to background contaminants using default parameters. A SAINTexpress FDR rate of ≤ 1% was selected as significant.

### Mass spectrometry data visualization and archiving

Functional enrichment analyses were performed with g:Profiler^102^ using default parameters. Dot plots and heat maps were generated using ProHits-viz (prohits-viz.org^103^). Interaction networks were generated using Cytoscape V3.5.1^104^, using the edge thickness to reflect each prey’s average spectral counts. Nodes were manually arranged into physical complexes. All MS files used in this study were deposited in MassIVE (http://massive.ucsd.edu) and assigned the MSV000095182. The username and password to access these files until publication is “asyn” at ftp://MSV000095182@massive.ucsd.edu.

### Western Blot

Whole cell lysates, from HEK293T-Rex cells stably expressing UltraID-LIPA constructs and exposed to blue light and with or without biotin for 30 min, were collected in 2X Laemmli lysis buffer (0.125 M Tris–HCl (pH 6.8), 20% glycerol, 0.2% 2-mercaptoethanol, 0.004% bromophenol blue, 4% SDS) after being washed 2 times with dPBS Ca+ Mg+. Samples were then prepared as previously described^22^. Briefly, samples were heated at 95 °C for 15 min to allow for DNA denaturation. Approximately 10 μl of the total protein fraction (corresponding to 35 μg of protein) was loaded per well in a 10% SDS-PAGE gel. The gels were run at 100 V for 90 min prior transfer of the protein to a nitrocellulose membrane using the Trans-Blot Turbo Transfer system (Bio-Rad, Hercules, CA, USA). The membranes were then incubated with a blocking buffer (PBS-Tween 0.1%, 3% Fish Gelatin) at room temperature for 1 h on a shaker set at a low rotation speed. Membranes were then incubated with an IRDye® 800CW Streptavidin (Licor, Cat. 926-32230) and finally washed 3 times 10 min with PBS-Tween 0.1%. Visualization and quantification were carried out with the LI-COR Odyssey scanner and software (LI-COR Lincoln, NE, USA).

### Coimmunoprecipitation assays (coIP)

HEK293T cells stably expressing the constructs lacking the UltraID (LIPA-α-syn, LIPA-empty and LIPA-TDP-43), seeded in 2*10-cm plates (per condition) were used for coIP experiments. Cells were exposed to blue light for a total of 1.5 h. HEK293T cells stably expressing the LIPA-α-syn construct not exposed to blue light were used as a negative control. One hour post blue light exposure or no light condition, media was removed, cells were washed once with warm dPBS Ca+ Mg+ to remove any free amines that might react with the crosslinker, and 50 μM disuccinimidyl glutarate (DSG) (Thermofisher, Cat. A35392) diluted in warm dPBS Ca+ Mg+ was added to the plates prior to incubation at 37 °C for 30 min under the blue light or in the dark. After incubation, dPBS containing DSG was removed, and cells were washed twice in cold dPBS Ca+ Mg+ to remove any remaining DSG. Whole cell lysates were scraped in lysis buffer (PBS-T 0.05%+ Protease inhibitor, Phosphatase inhibitor II, Phosphatase inhibitor III, and PMSF). The lysates were kept on ice for 5 min. Once the cells were swollen, cells were lysed through gentle shearing with a syringe equipped with a 27 G gauge needle (Fisher Scientific, 14-826-87). The progress of cell lysis was monitored by phase contrast microscopy to ensure sufficient lysis, using 25 syringe cycles. Samples were sequentially centrifuged at 4 °C (500 x g for 5 min and 1,000 x g for 10 min). The supernatant was collected and 10% of the volume was kept as input sample and diluted in NuPage Sample Buffer (4X) containing lithium dodecyl sulfate (LDS) (Thermofisher, Cat. NP0007). The remaining supernatant was then used for coIP.

Immunoprecipitation was performed using Dynabeads Protein G (Thermofisher, Cat. 10004D) following the manufacturer’s instructions. First, a volume of 50 μL (per 500 μL of sample) of dynabeads was transferred to clean microcentrifuge tubes. Tubes were placed on the magnet to separate the beads from the solution. 2 µg of the mCherry antibody diluted in 200 μL PBS-T 0.05%, was added to the magnetic beads. The beads-antibody complex was incubated with rotation for 30 min at room temperature. The beads were then washed in PBS-T 0.05% 3 times by gentle pipetting. After the last wash, samples were added to the magnetic bead-antibody complex, the samples were incubated with rotation for 16 h at 4°C to allow the antigen to bind to the magnetic bead-antibody complex. After the incubation time, the tube with the samples were placed on the magnet, and supernatant was removed. The magnetic bead-antibody-sample complex was washed 3 times by gentle pipetting using 200 μL of PBS-T 0.05%. The magnetic bead-antibody-sample complex was resuspended in 100 μL of PBS-T 0.05% and transferred to a clean tube. To elute the samples, 20 μL of elution Buffer (50 mM glycine at pH 2.8) was used, and 10 μL of premixed NuPAGE LDS Sample Buffer and 2-Mercaptoethanol (5%, final concentration). The samples were then heated for 10 min at 70°C. Samples were then subjected to immunoblotting as described in the Western Blot protocol.

### Immunocytochemistry

HEK293T cells, grown on poly-L-lysine (Sciencell, Cat. 0413) coated #1.5 8-mm glass coverslips (Electron Microscopy Sciences, Cat. 72296-08), were fixed with 4% (wt/vol) paraformaldehyde (PFA) (Electron Microscopy Sciences, Cat. # 19210) and 3% (wt/vol) sucrose (Sigma-Aldrich, Cat. S9378-1KG) for 15 min at room temperature. After three 5-min washes with 1× PBS, the cells were permeabilized with 0.3% (vol/vol) Triton X-100 in 1× PBS for 5 min. Cells were then rinsed 3 times for 5 min each with 1× PBS to remove remaining Triton, and further incubated with a blocking buffer constituted of 5% (vol/vol) normal goat serum (NGS) (Thermo Scientific, cat. 16210064), 1% (wt/vol) BSA (BioShop, Cat. ALB001.500) and 0.1% (vol/vol) saponin (Sigma-Aldrich, Cat. S4521-10G) diluted in 1× PBS) for 1 h at room temperature. The cells were subsequently incubated with primary antibodies (WDR44 1:200; VPS13C 1:100; BLTP3A 1:100; IGFR2 1:100 or pS129 1:2000) (referenced in the key resources table), diluted in the blocking solution and incubated overnight at 4°C. After three 5-min washes with fresh blocking buffer, the cells were incubated with secondary antibodies Alexa Fluor 488 (Invitrogen, Cat. A11008) diluted at 1:800 for 1 h at room temperature. After three 5-min washes with 1× PBS, the cells were stained with DAPI (4’,6-diamidino-2-phenylindole) (Thermo Fisher Scientific, Cat. 1729803) diluted at 1:5000 in 1× PBS for 2 min and then washed twice with 1× PBS. Prior to mounting, the cells were quickly rinsed with mQH2O to remove remaining salts. Finally, the coverslips were mounted on slides using 3.5 μL of Fluoromount-G with a refractive index of ∼1.4 (Thermo Fisher, Cat. 5018788). Slides were dried overnight at room temperature in the dark prior to imaging.

### Confocal imaging and aggregates segmentation/quantification

The following imaging parameters are reported following the recommendations by Llopis *et al*.^105^. Confocal images were acquired on a Zeiss LSM800 upright microscope, with a motorized stage (WSB Piezo Drive CAN) and equipped with a beam splitter for 4 non-tunable solid-state lasers: 405 nm – 488 nm – 591 nm – 640 nm. Images were acquired using the 405 nm for nuclear staining (DAPI), 488 nm for the specific markers (Alexa Fluor 488) and the 581 nm was used for the LIPA constructs (mCherry). Laser intensities and gain were modulated to optimize the signal-to-noise ratio and avoid saturation using the range indicator mode: 0.1% (405 nm), 0.04 to 0.1% (488 nm and 591 nm) while the digital gain was kept between 600 V (for conditions with bright and dense aggregates, avoiding saturation of the signal) and 750 V (mostly for the conditions without any aggregates, to emphasize the diffuse cytosolic signal). Point laser scanning was performed using the LSM800 scan head galvo scanning mirrors with a unidirectional scan set at 900 Hz with 2-line averages. For all the images, a 40x objective Plan-Apochromat 40x Oil DIC (UV) VIS-IR M27/1.40 NA (Zeiss, Cat. 420762-9900-000) was used with an immersion oil Type LDF (Cargille, cat. #16919-16) (refractive index 1.5180 +/- 0.0002). The wavelength emitted by the fluorophores was collected by a GaAsP-PMT1 photomultiplier tube set to capture the fluorescence emitted ranging from 600 to 700 nm (mCherry), 400 to 575 nm (Alexa Fluor 488) and 400 to 500 nm (DAPI), with a pinhole size ranging from 0.15 AU to 0.24 AU. Sequential scanning between each z-stack was performed, acquiring the emitted fluorescence from the longest wavelengths (mCherry) to the shortest (DAPI). Z-stacks were acquired applying Nyquist sampling rates to reach a voxel size of 0.5 micron. A numeric zoom of x1 or x3 was applied using an Axiocam 506 mounted to the phototube with a Zeiss 1.0x C Mount Camera Adapter "60N" to achieve a resolution of 512x512 pixels or 1024x1024 pixels to reach a pixel size of 156 nm. All the acquisition parameters were set on Zen 3.9.0. Image analysis, color balance, contrast and brightness were adjusted using the open-source software Fiji ImageJ (https://imagej.nih.gov/ij/). Aggregates were segmented directly from the z-stack using the plugin Distance Analysis (DiAna)^106^, to analyze the average volume of the aggregates. For an optimal segmentation, the detection and gaussian segmentation threshold were adjusted for each image in the mCherry channel using a threshold between 50 and 70 excluding the values below 2 and above 90000 pixels that were considered as a signal background. The analysis of the average number of aggregates per cell was performed manually using the manual counter tool incorporated in ImageJ. An aggregate was considered as a unique entity if it could be delineated by a continuous line. Two close aggregates were considered as different entities if each could be delineated by a continuous trace.

### Statistical analysis

Quantification of LIPA aggregates: aggregate quantifications were conducted in at least 3 independent experiments and were further analyzed using Prism v.9 (GraphPad, La Jolla, CA, USA). Statistical analysis: abnormality test was performed. If the normality test was positive, we then performed a parametric test (Student’s t-test). If the normality test was negative, in cases where we analyzed more than two conditions, a one-way ANOVA test was performed. All values were expressed as means ± SEM. A *p*-value < 0.05 was required to reject the null hypothesis.

Experimental design for MS experiments: for each bait, 3 biological replicates were independently processed. Each batch of samples processed included negative controls, which were grown in parallel to the bait samples and treated in the same manner. We used UltraID-LIPA-empty cells to model the background protein hits in our experiments. To minimize sample carry-over issues during liquid chromatography, extensive washes were performed between samples and the order of sample acquisition was randomized.

---

**Figure S1:**
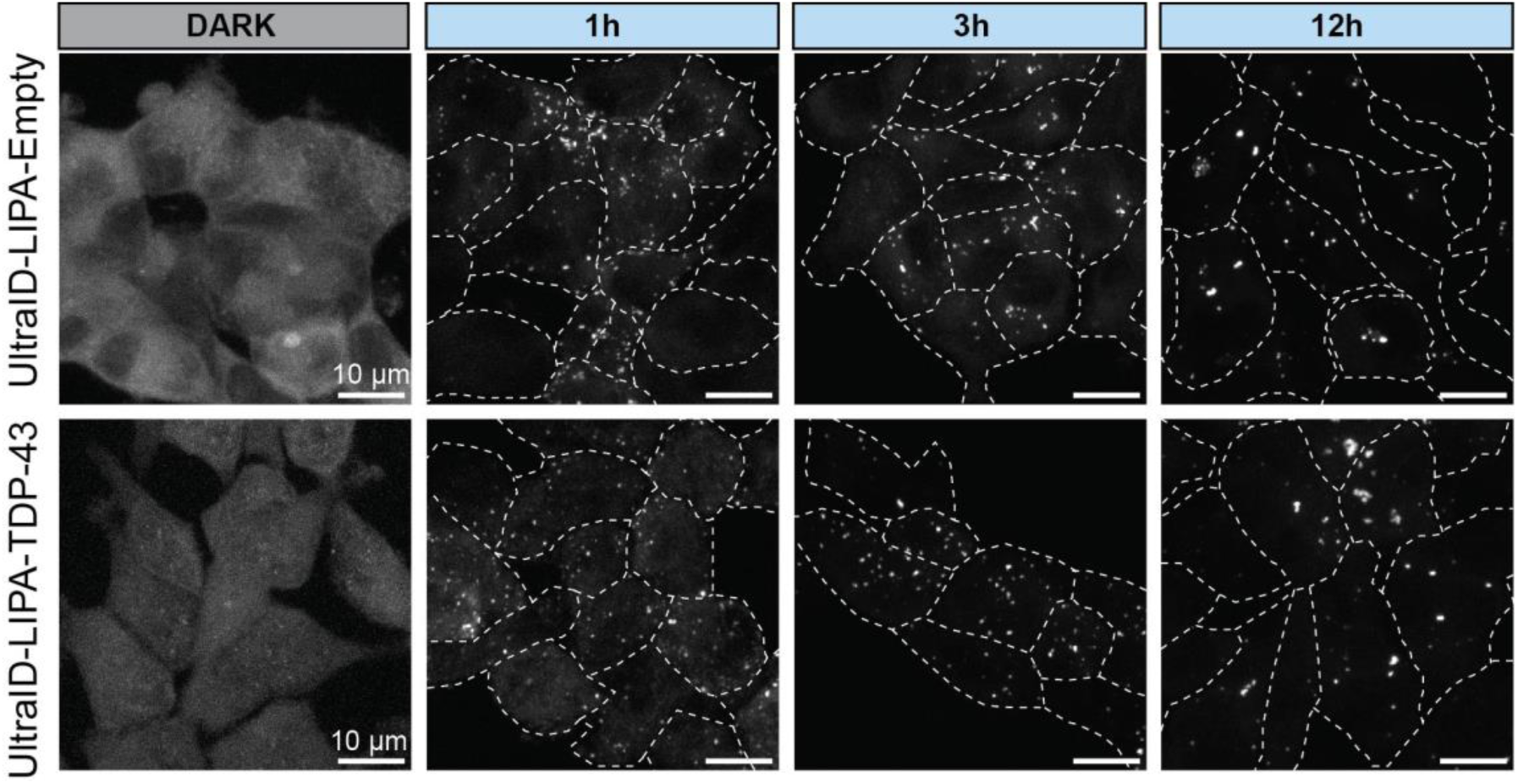
UltraID doesn’t affect LIPA-Empty or LIPA-TDP-43 aggregation propensity. Confocal maximum intensity projections images of Flp-In T-REx HEK293T cells stably expressing UltraID-LIPA-α-syn construct. Cells were exposed to blue light for 1h, 3h and 12h to induce the aggregation. Scale bar = 10 µm.

**Figure S2.**
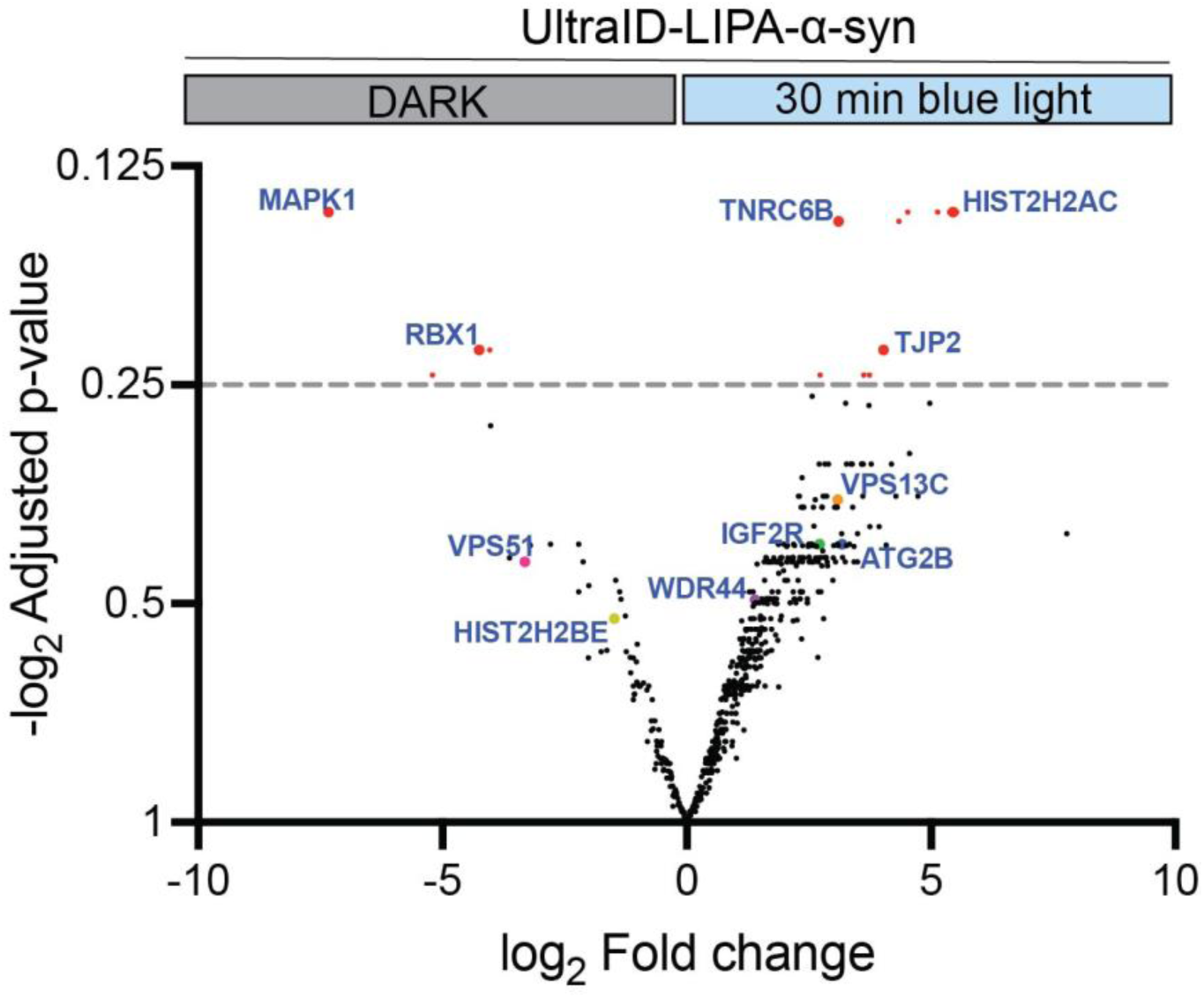
Proteome analysis reveals significant enrichment in proteins in the α-syn oligomers induced by the LIPA system: Volcano plot representing mass spectrometry analysis of the protein content of all the replicates (n=3) for the conditions exposed or not to the blue light. Each of the 684 proteins identified in the present proteomic study is plotted as a circle positioned upon the fold change (log_2_ Fold change) and statistical significance (−log_2_ Adjusted *p*-value) of its enrichment in α-syn monomers (DARK, left side) and α-syn oligomers (LIGHT, right side). A total of 567 proteins were found to be significantly enriched in the oligomers group and 117 proteins were found to be enriched in the monomers group. Proteins of interest are highlighted in bold with larger dots matching the corresponding text color. Red dots indicate proteins that are highly significantly enriched on both sides (above 0.25 −log_2_ Adjusted *p*-value).

**Figure S3:**
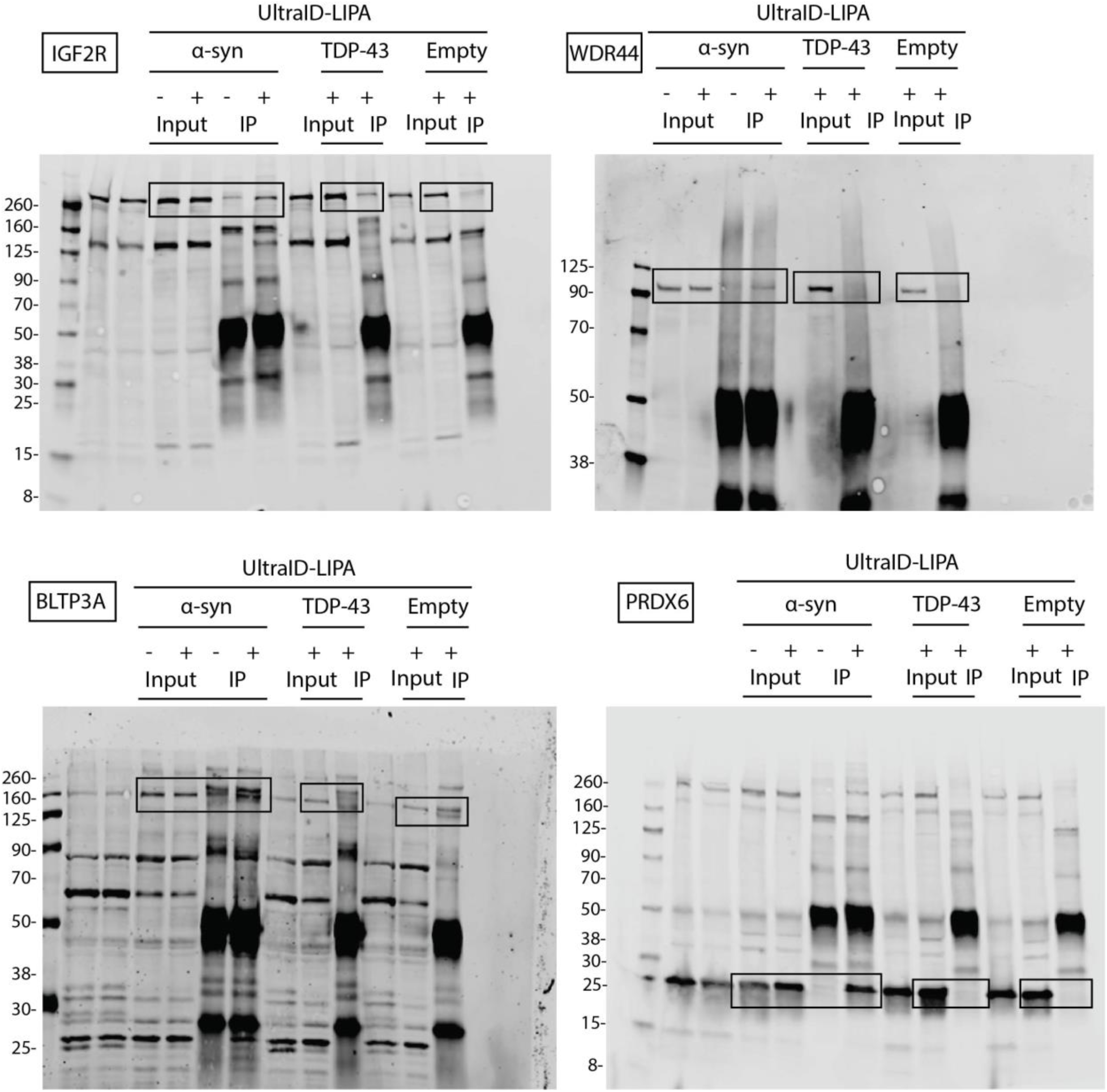
Uncropped membrane scans for all presented Western blot in Fig. 4A.

